# Fluorescent Sensors for Imaging Interstitial Calcium

**DOI:** 10.1101/2023.03.23.533956

**Authors:** Ariel A. Valiente-Gabioud, Inés Garteizgogeascoa Suñer, Agata Idziak, Arne Fabritius, Julie Angibaud, Jérome Basquin, U. Valentin Nägerl, Sumeet Pal Singh, Oliver Griesbeck

## Abstract

Calcium in interstitial fluids is central to systemic physiology and a crucial ion pool for entry into cells through numerous plasma membrane channels. Its study has been limited by the lack of methods that allow monitoring in tight inter-cell spaces at high spatio-temporal resolution. We engineered high performance ultra-low affinity genetically encoded calcium biosensors named GreenT-ECs. GreenT-ECs combine large fluorescence changes upon calcium binding and binding affinities (K_D_) ranging from 0.8 mM to 2.9 mM, making them uniquely tuned to calcium concentrations in extracellular organismal fluids. We validated GreenT-ECs in rodent hippocampal neurons and transgenic zebrafish *in vivo*, where the sensors enabled monitoring homeostatic regulation of tissue interstitial calcium. GreenT-ECs may become useful for recording very large calcium transients and for imaging calcium homeostasis in inter-cell structures in live tissues and organisms.

## Introduction

Calcium acts as a crucial second messenger within cells, involved in regulating a plethora of cellular processes. The complexities of these cellular signaling mechanisms, their spatiotemporal dynamics and the many channels, buffers, pumps and downstream effector enzymes involved have been studied extensively^1, 2^. In contrast, calcium dynamics and regulation in interstitial fluids have been far less studied. Free ionic calcium in these fluids forms a critical reservoir for entry into cells through several classes of plasma-membrane calcium channels. It is in equilibrium with bound calcium in proteins, extracellular matrix components and is present in high amounts in bone and teeth. Multicellular organisms have developed means to tightly control and regulate its concentrations in circulating fluids. A central role here is played by the Calcium Sensing Receptor (CaSR), a G-protein coupled receptor that, among other ligands, senses extracellular calcium and transduces intracellular signaling^3, 4^. Its expression in the parathyroid gland, the secretion of parathyroid hormone by dedicated cells in the parathyroid gland, and the coupling to active vitamin D^5-7^ signaling are key components of a central regulatory system that adjusts free calcium in body fluids to a narrow concentration range^7^. While central mechanisms of calcium homeostasis in interstitial fluids appear well understood, less is known about its regulation in extended and narrow inter-cell spaces that may be finely spun and far away from known homeostatic regulatory centers. The use of ion-selective electrodes, magnetic nanoparticles and micro-dialysis has opened a window into complex dynamics and modes of its regulation in extracellular volumes^8, 9^, but typically these techniques suffer from low spatiotemporal resolution. Imaging of fluorescent calcium indicators, in combination with advanced microscopy, offers excellent spatiotemporal resolution^10^. Genetically encoded calcium indicator variants^11^ allow precise localization in tissues and at subcellular sites of interest. Unfortunately, the focus of sensor development lay almost exclusively on applications within cells or cellular organelles, resulting in excellent sensors with high (nM) affinities for calcium. Sensors with intermediate to low affinity were engineered to image calcium in cellular organelles where free calcium concentrations may reach levels of several hundred micromolar^12-15^. However, even lower affinities are needed to measure larger calcium fluxes and to match concentrations encountered in various body fluids.

## Results

### Development of GreenT-ECs

For imaging calcium in body fluids and inter-cell spaces, we engineered a new class of low affinity genetically encoded calcium sensors (Fig. 1). Our design was inspired by the molecular architecture of Camgaroo^16, 17^. As calcium sensing moiety we employed a minimal calcium binding domain derived from Troponin C (TnC)^18^ and used it to replace Tryptophan 148 within the bright green fluorescent protein mNeonGreen^19^. A similar design for a high affinity sensor had been presented earlier^20^. Compared to the GCaMP design^21, 22^ prominent in high affinity biosensors, our design yielded sensors with fewer calcium binding sites per indicator (2 vs 4) and conserved original N-and C-termini of mNeonGreen for fusion of subcellular targeting motifs. Due to the bright green fluorescence in bound state, the insertion of the TnC fragment and the tuning of affinities for extracellular applications we named the sensors GreenT-ECs. To elucidate more details on sensor function and for guidance for our protein engineering efforts we solved a high-resolution crystal structure of an early variant of the sensor in the calcium bound state to 1.3 A° resolution (Fig. 1A, Table S1). It revealed a structure in which the ß-barrel of mNeonGreen is opened to harbor the minimal calcium-binding domain of TnC. Within the TnC domain the two EF-hand loops with two bound calcium ions and key regions linking the TnC domain to mNeonGreen are visible. Using structure guided optimization and extended iterative directed evolution (Fig. 1B) we were able to dramatically enhance fluorescence changes and response behavior of the sensors. Fluorescence changes finally were about 60-fold in response to calcium binding and basal fluorescence in the absence of calcium was close to detection limits (Fig 1C) (Table 1). The excitation and emission maxima were 504 nm and 515 nm, respectively. There was no detectable interference by magnesium (Fig. 1D). The binding affinity was drastically reduced by inserting either an additional aspartic acid after position 168 (GreenT-EC) or a serine after position 205 (GreenT-EC.b) within the EF-hand calcium chelating loops. Additionally, GreenT-EC.b, bearing the extra mutation T261I (GreenT-EC.c) presented the lowest calcium-binding affinity (Fig. S1, Table 1). Altogether, these mutations allowed tuning the sensors to the low affinities necessary for use in interstitial fluids, without compromising the large fluorescence changes achieved. *In vitro* affinity titrations of purified recombinant proteins revealed sensors with Kds in the millimolar range. A comprehensive *in vitro* analysis of the spectroscopic properties of the sensors is available in Figure 2. GreenT-ECs could be localized to the surface of mammalian cells by fusion of a number of appropriate targeting motifs (Fig. S2). The fluorescence properties of GreenT-ECs facilitated optimization of this process enormously, as the indicators, when expressed in the cytosol, are almost non-fluorescent while brightly fluorescent when exposed to millimolar calcium concentrations found in extracellular liquids or buffers (Fig. S3). This allowed achieving an exquisite signal contrast between bright fluorescence of surface exposed indicators versus extremely dim fluorescence of residual indicators inside cells within the cytosol or *en route* to the plasma membrane. Conceivably, dynamic probing of surface exposure or secretion of proteins might be an interesting secondary use of GreenT-ECs. A reference protein could be fused to the cytosolic C-terminal end of the sensor, facilitating calibration measurements. Among several proteins tested (Fig. S4), the red fluorescent protein mCyRFP1^23^ was a useful candidate that allowed convenient co-excitation of the two fluorophores with a single laser wavelength. Referenced GreenT-ECs could be titrated on the surface of HEK293 cells by sequentially increasing the free calcium concentration in extracellular buffers (Fig. 1H). These titrations yielded higher observed affinities than the measurements performed with purified recombinant protein. The Kds for calcium were 1.3 mM (GreenT-EC), 0.8 mM (GreenT-EC.b) and 2.9 mM (GreenT-EC.c) (Fig 1H) (Table 1).

**Figure 1:**
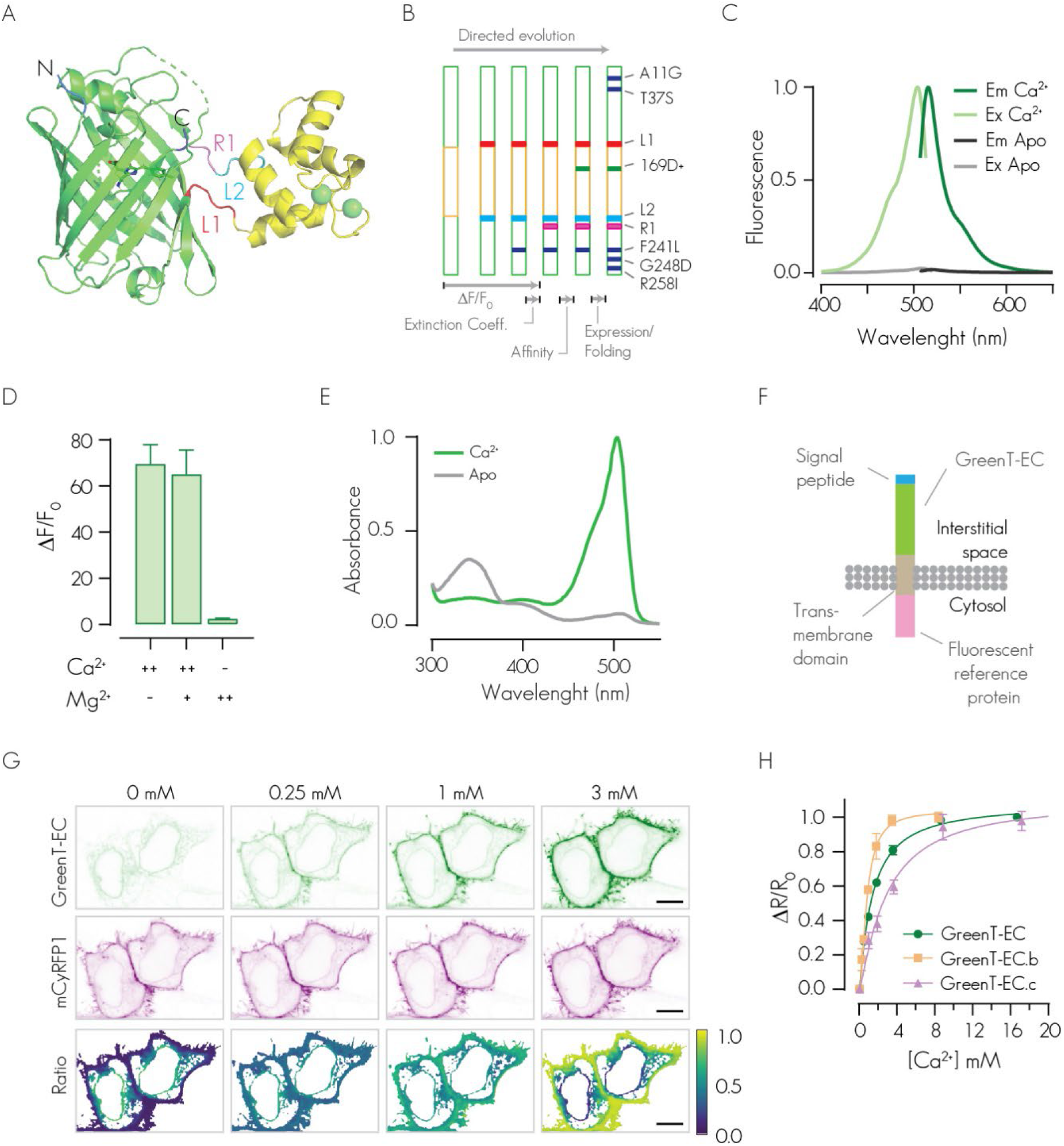
Development of GreenT-ECs. **A**) Crystal structure of a calcium bound GreenT-EC intermediate variant (NRS F241Y) generated during the process of directed evolution. The minimal calcium-binding domain derived from Troponin C (yellow) was inserted into the fluorescent protein mNeonGreen (green). Spheres (green) indicate two bound calcium ions. Linkers (L1 and L2) as well as a particularly crucial region (R1) for engineering are highlighted. N and C mark the N-and C-terminus of the protein. **B**) Iterative cycles of directed evolution were performed to optimize GreenT-ECs. The scheme represents the indicator (green: mNeonGreen moiety, yellow: TnC minimal domain). The major evolutionary steps illustrate sites of amino changes that lead to functional improvements. L1, L2: mutated linker amino acids. R1: stretch of three amino acids comprising residues 225-227 reaching into the ß-barrel. Together with linker mutations, this sensitive area fundamentally affected fluorescence change, brightness and off-kinetic of variants. Individual amino acid changes affecting folding and expression are indicated (blue). **C**) Excitation and emission spectrum of GreenT-EC at 0 mM (gray) and 30 mM (green) Ca^2+^. The excitation and emission maxima are 504 nm and 515 nm, respectively. **D**) Maximal fluorescence change of GreenT-EC in response to calcium and/or magnesium (++ is 100 mM, + is 5 mM, - is 0 mM). **E**) Absorbance changes of GreenT-ECs in the absence (grey) and presence (green) of calcium. The major absorbance peak maximum was at 504 nm. Minor absorbance peaks at 350 nm and 400 nm in the absence of calcium corresponded to dark forms of the chromophore. **F**) Cell surface display of GreenT-EC. An effective way consisted of fusing an N-terminal signal peptide to GreenT-EC and a C-terminal membrane anchoring domain (from PDGF receptor). The fluorescent reference protein mCyRFP1 was added to the C-terminus (intracellular). **G**) Cell surface titration of calcium affinities of GreenT-ECs. HeLa cells expressing surface delivered GreenT-ECs were exposed to increasing calcium concentrations. Green (GreenT-EC) and red (mCyRFP1) emissions are detected simultaneously and are presented in the upper and middle panels respectively. Ratiometric (GreenT-EC/mCyRFP1) images are presented in the bottom panel. Scale bar, 10 mm. **H)** Summary of GreenT-EC/mCyRFP1 titrations on the surface of HEK293 cells. The mean normalized response ±SEM is plotted (Three independent experiments per variant, 8-10 cells per experiment).

**Figure 2:**
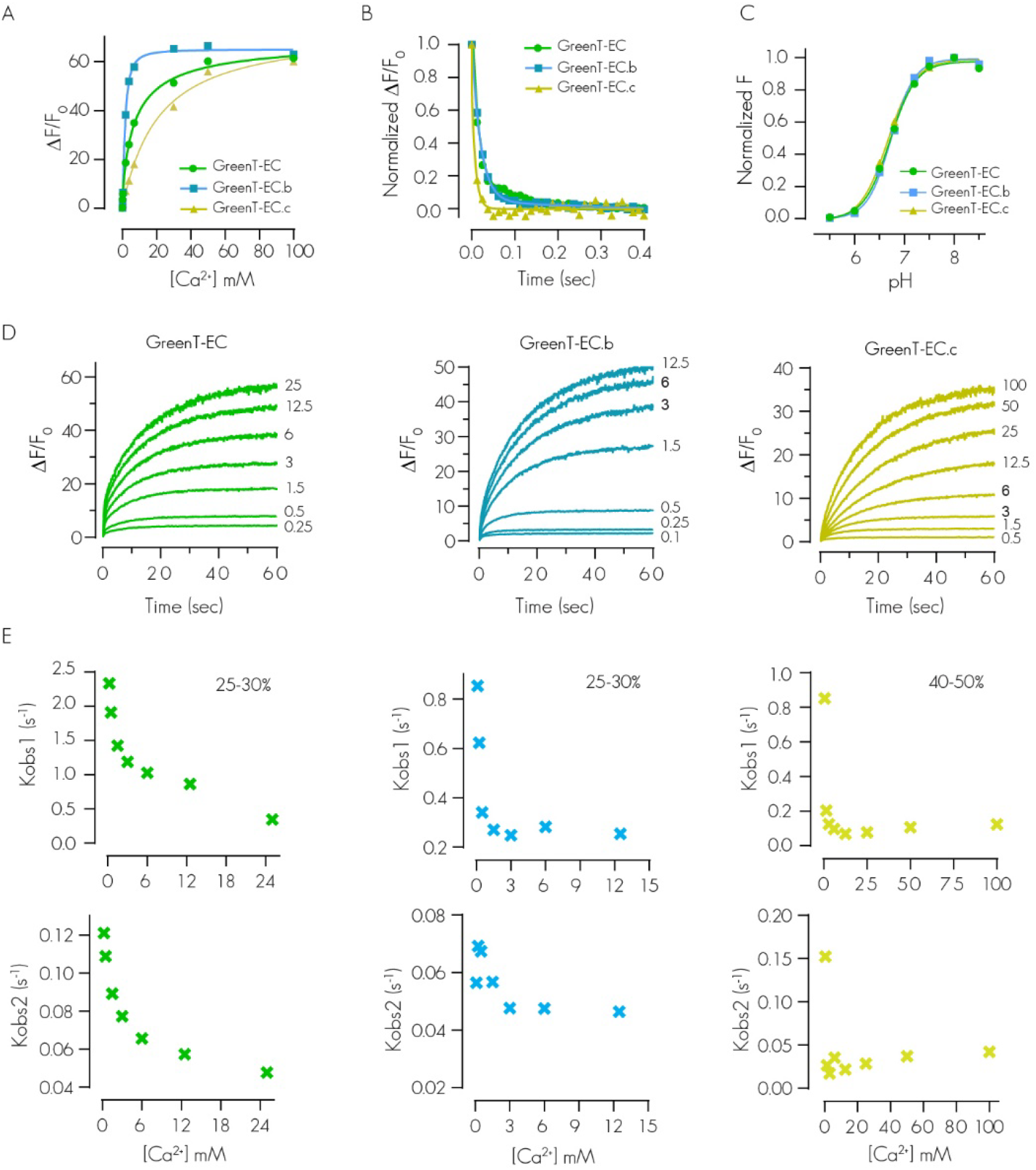
In vitro spectroscopy of GreenT-ECs. **A**) In vitro affinity titration curves of recombinant purified GreenT-EC and two variants with higher (GreenT-EC.b) and lower (GreenT-EC.c) affinity. **B**) Off-rate kinetics of the three variants of the sensor. The proteins were prepared in MOPS 30 mM, KCl 100 mM, Mg 2.5 mM, Ca2+ 100 mM and rapidly mixed with BAPTA 200 mM. **C**) pKa of the three sensors studied in this article. Measurements were obtained by preparing pH solutions in MOPS/MES containing 100 mM Ca2+. **D**) On-rate kinetics of GreenT-EC (green), GreenT-ECb (blue) and lower affinity GreenT-ECc (yellow) were measured by rapidly mixing solutions of protein in MOPS 30 mM, KCl 100 mM, Mg 1 mM with MOPS 30 mM, KCl 100 mM, Mg2+ 1 mM containing increasing concentration of Ca2+. **E**) A double exponential was used for the fittins for each variant and the obtained Kon values were plotted against the calcium concentration. Kobs1 corresponds to the first step of the rate, representing 25-30 % of the total response. Kobs2 corresponds to the slower process and accounts for the higher percentage of the total response. In all cases, a decrease in the Kobs is observed at increasing concentrations of calcium. This results suggest a mechanism by which there is an equilibrium between at least two species of the sensor, and only one can progress to a fluorescent species upon calcium binding.

**Table 1:** Spectroscopic properties of GreenT-ECs. ΔF/F_0_, fluorescence change upon calcium binding; EC, extinction coefficient of the bound state; K_D_, dissociation constant; QY, quantum yield of the bound state; n_H_, Hill coefficient; Koff, off-rate kinetics; Kobs1 and Kobs2, the observed on-rate kinetics of the two-phase association process calculated at calcium saturation. The percentage of the response obtained during the fast phase is indicated. ΔR/R_0_, ratio change on HEK293 cells using mCyRFP1-referenced sensors.

| Variant | In vitro properties |  |  |  |  |  |  |  |  |  | Cell surface titration |  |  |
| --- | --- | --- | --- | --- | --- | --- | --- | --- | --- | --- | --- | --- | --- |
| | $\Delta F/F_0$ | EC<br>(mM<sup>-1</sup><br>cm<sup>-1</sup>) | QY | QY*EC | K<sub>D</sub><br>(mM) | n<sub>H</sub> | K<sub>off</sub><br>(s<sup>-1</sup>) | K<sub>obs1</sub><br>(s<sup>-1</sup>) | K<sub>obs2</sub><br>(s<sup>-1</sup>) | pKa | $\Delta R/R_0$ | k<sub>d</sub><br>(mM) | n<sub>H</sub> |
| GreenT-EC | 61 | 79 | 0.72 | 57 | 7.1<br>± 1.2 | 0.8<br>± 0.1 | 33 | 0.35<br>(25%) | 0.05 | 6.7 | 10<br>± 3 | 1.3<br>± 0.2 | 1.1<br>± 0.2 |
| GreenT-EC.b | 58 | 81 | 0.74 | 60 | 1.6<br>± 0.1 | 1.5<br>± 0.1 | 50 | 0.25<br>(25%) | 0.05 | 6.7 | 8<br>± 2 | 0.8<br>± 0.1 | 1.8<br>± 0.6 |
| GreenT-EC.c | 56 | 80 | 0.72 | 58 | 20<br>± 12 | 1.0<br>± 0.2 | 50 | 0.12<br>(40%) | 0.04 | 6.7 | 10<br>± 2 | 2.9<br>± 1.9 | 1.2<br>± 0.6 |

### Validation of GreenT-EC in rodent hippocampus

We next validated sensors in cultured rat hippocampal neurons and mouse hippocampal slices (Fig. 3). For these experiments, we anchored GreenT-EC to the neuronal surface with a GPI-anchor (Fig. 3A)^24^. Both in transfected hippocampal neurons in culture (Figure S5) and in organotypic slices after AAV (Adeno-Associated Virus) delivery of GreenT-EC (Fig. 3B) we saw bright and specific surface labeling of transfected cells using confocal microscopy and super-resolution STED (stimulated emission depletion) respectively. Fluorescence was resistant to photobleaching, allowing for repetitive imaging of the same cells (data not shown). Moreover, fluorescence intensity dynamically changed when perfusing neurons with buffers containing 0 mM, 1.5 mM and 8 mM free calcium (Fig. S5, Fig. 3C). This confirmed that the indicator affinity was well placed to report both potential decreases and increases in interstitial calcium. In addition, in organotypic brain slices kept in buffered medium at 1.5 mM Ca^2+^, GreenT-EC signals dynamically increased when solutions containing 8 mM free calcium were locally puff-applied using a picospritzer (Fig. 3D-E). We next evoked neuronal activity by electrically stimulating the Schaffer collaterals and recorded GreenT-EC responses in the CA1 projection area using two-photon microscopy (Fig. 3F). We detected transient increases in GreenT-EC signals after stimulation. Similarly, spontaneous neuronal activity elicited increases in GreenT-EC signals (data not shown). We set out to explore the origins of these calcium rises in hippocampus using pharmacology. A combination of sodium orthovanadate (SOV, 5 µM) and benzamil hydrochlorate hydrate (BHH, 50 µM), inhibitors of the plasma membrane calcium ATPase and the Na^+^/Ca^2+^ exchanger, essentially blocked the increases in interstitial calcium after Schaffer collateral stimulation (Fig. 3G). Interestingly, recorded GreenT-EC signals were larger and longer lasting in the deeper layers of slices compared to more superficial layers (Fig. 3H) where extruded calcium may be dissipating more easily. Possibly, the compactness and size of the interstitial space could have an effect on shaping these signals.

**Figure 3:**
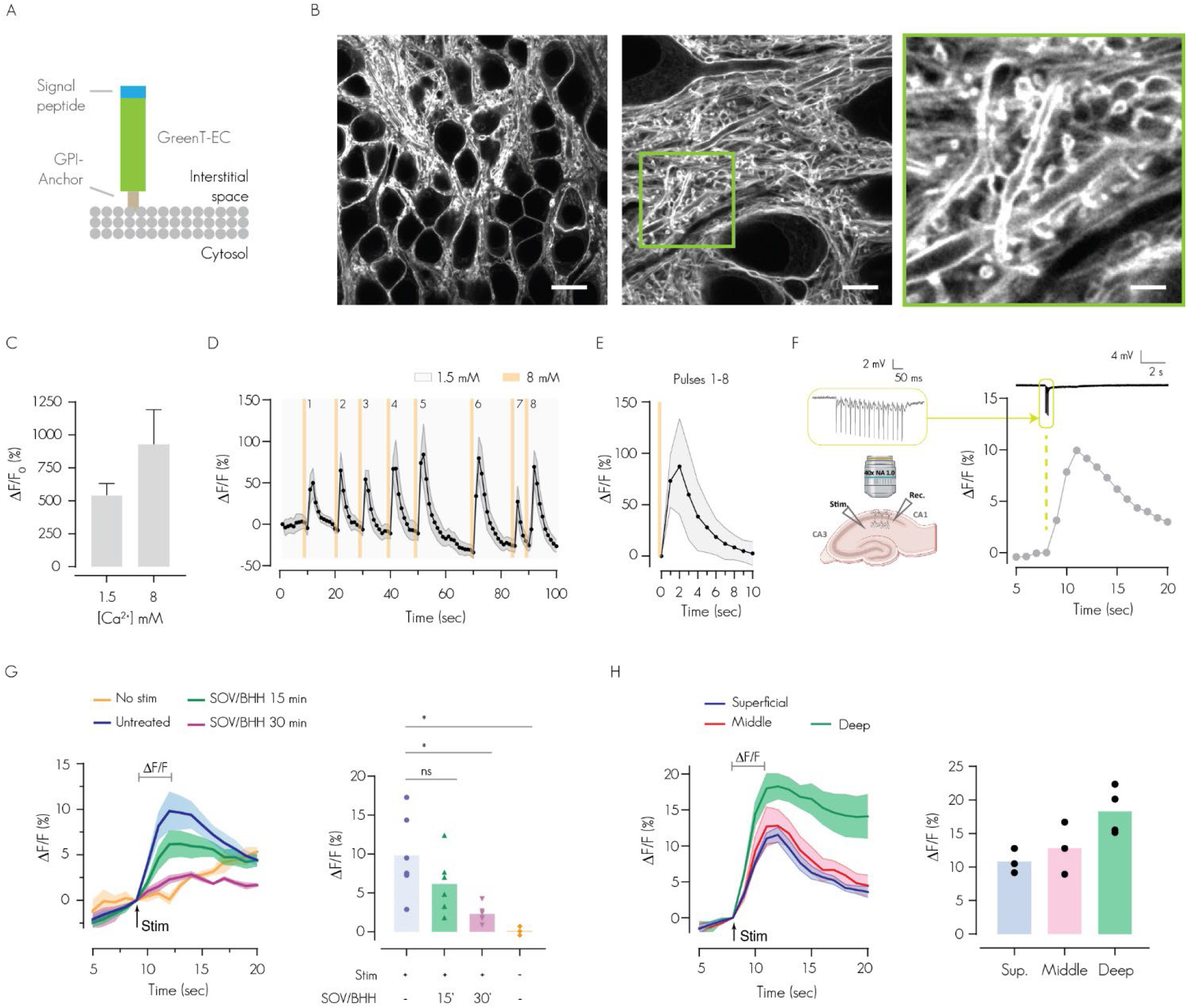
Validation of GreenT-EC in rodent hippocampus. **A**) Scheme of surface targeted GreenT-EC (green) by means of an N-terminal signal peptide (blue) and a C-terminal GPI anchor (brown). **B**) STED images of AAV-GreenT-EC transfected neurons in hippocampal area CA1 of organotypic slice cultures (7 day post infection). Scale bars: 10 μm, 5 μm, 2 μm, respectively. **C**) Maximal fluorescence changes in transfected dissociated hippocampal neurons (N = 4) from zero calcium to 1.5 mM and 8 mM extracellular calcium. **D**) Dynamic GreenT-EC fluorescence changes upon repetitive brief (500 ms each) high-calcium (8 mM) solution puffs. Buffer calcium was at 1.5 mM. Orange lines indicate timing of the puffs. **E**) Averaged response curves to the calcium injections from experiment D. **F**) Electrical stimulation of the hippocampal Schaffer collateral pathway was used to elicit GreenT-EC responses monitored by 2-photon microscopy. A GreenT-EC response in hippocampal CA1 is shown. **G**) Activity evoked hippocampal CA1 GreenT-EC signals and their block by a combination of sodium ortho-vanadate (SOV, 5 µM) and benzamyl hydrochlorate hydrate (BHH, 50 µM) at 15 or 30 minutes after drug infusion. (**H**) Activity-induced hippocampal GreenT-EC fluorescence signals as a function of imaging depth within the organotypic slice. 3-6 experimental replicates were evaluated for each condition. One-way ANOVA followed by a Dunnett’s test was used to evaluate differences among control and treated groups (* p < 0.05).

### *In vivo* imaging of calcium homeostasis in transgenic zebrafish

To establish a model of body fluid ionic homeostasis, we next generated transgenic zebrafish that stably expressed mCyRFP1-referenced GreenT-EC semi-ubiquitously using the 10 kb *actb2* promoter^25^ (Fig. 4). The transgenic fish displayed green and red fluorescence (Fig. 4A-C, E), exquisitely outlining various cell types such as muscle cells, epithelial cells or vacuolar cells of the notochord in finest detail and without signs of aggregation of the indicator (Fig. 4C). For volumetric imaging the regulation of live tissue interstitial calcium, we performed confocal imaging of the dorsal fin fold and posterior notochord after various types of treatment (Fig. 4D). To validate the sensor functionality, we initially incubated 4-day post fertilization (dpf) zebrafish larvae acutely for 10 min in 3 mM EGTA. *In vivo* confocal imaging revealed a drastic loss of green fluorescence throughout tissue volumes, while red cytosolic mCyRFP1 fluorescence remained stable (Fig. 4E, F). Longer treatments with EGTA lead to considerable tissue disintegration. We next applied a model of hypocalcemia. In this, larvae are transferred to low-calcium water (0.03 mM), which is thought to induce a feed-back loop that increases the capacity of the animal to absorb calcium via increased calcium sensing receptor (CaSR) expression and proliferation of CaSR-expressing cells^26, 27^. We incubated transgenic larvae from 2 dpf on in low calcium water enriched with the CaSR inhibitor Calhex231 (5 and 10 µM)^28^ and imaged the fin fold at 4 dpf using *in vivo* confocal microscopy. Remarkably, treatment with Calhex231 led to a significant and dose dependent decrease in interstitial calcium, although the effect was tissue-specific. We saw less pronounced effects in muscle tissue, possibly hinting at a less prominent role for CaSR in calcium homeostasis in muscle or a lower diffusion rate of interstitial calcium in muscle. (Fig. 4E, G). Under similar experimental conditions and incubation periods Calcitriol (2.5 µM), the active form of Vitamin D and a hypercalcemic hormone in zebrafish^29^, did not lead to detectable whole tissue changes of fluid calcium levels. Higher concentrations of Calcitriol led to animal death. Finally, we were interested in exploring the natural capacity of larval zebrafish for maintaining calcium homeostasis under conditions of large variations in calcium concentrations in the external environment (Fig. 4I). We raised fish from birth in embryonic medium with large concentration differences in calcium (0.03 mM – 10 mM, pH 7.4). When these fish were imaged at 2, 4 or 6 dpf no significant differences in body fluid calcium could be detected, demonstrating the remarkable ability of homeostatic mechanisms to clamp interstitial calcium levels to the physiological range. It is worth to highlight the stability of GreenT-EC/mCyRFP1 across animals considering that experiments were acquired over repeated sessions spread out over long periods of time.

**Figure 4:**
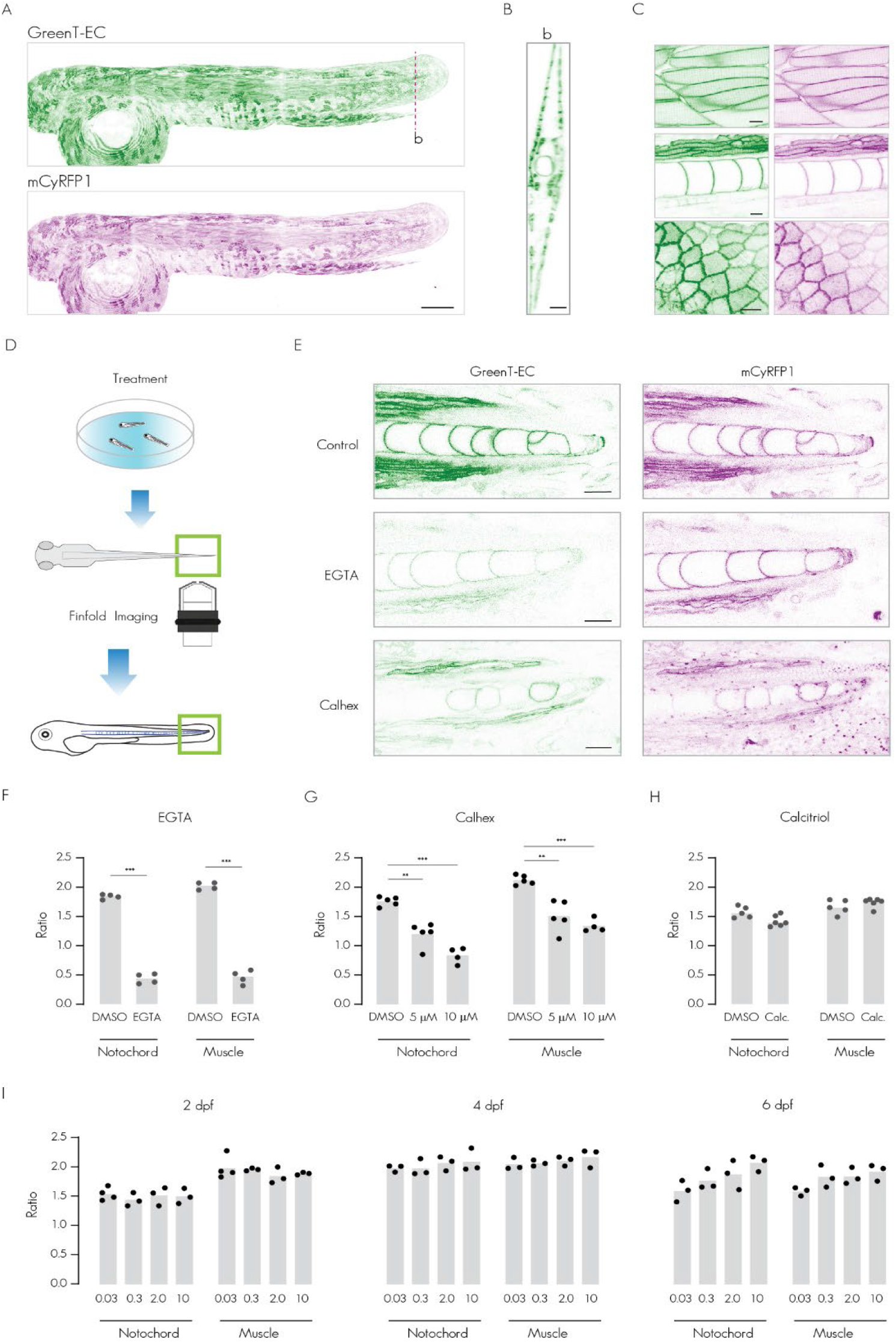
In vivo imaging of calcium homeostasis in transgenic zebrafish. **A**) Maximum intensity Z-projection of a confocal stack from a 2 dpf transgenic zebrafish expressing mCyRFP1-referenced GreenT-EC under the control of the β-actin promoter. GreenT-EC (green) as well as mCyRFP1 emission channels (magenta) are displayed. Scale bar, 200 μm. **B**) Representative orthogonal view of the animal model in the fin fold. Scale bar, 20 μm. **C**) Representative images of skeletal muscle cells (upper panel), vacuolar cells of the notochord and ventral/dorsal muscle (middle panel) and epithelial cells (bottom panel) in the fin fold area of zebrafish. Note the excellent membrane translocation of the sensor in all tissues, the lack of aggregates and the low background coming from intracellular structures. Scale bar, 10 µm. **D**) Experimental strategy used for studying calcium homeostasis in transgenic GreenT-EC zebrafish larvae. After acute or chronic treatment fish are mounted in low melting agarose oriented as depicted and the posterior fin fold is imaged. **E**) Confocal two channel (GreenT-EC/mCyRFP1) images of the fin fold of 4 dpf zebrafish larvae after various treatments. Top: fish larvae under control conditions. Middle: after 10 min treatment with EGTA (3 mM). Bottom: treatment with Calhex231 (10 µM) from 2 dpf to 4 dpf. Scale bar, 20 µm. **F**) Quantification of the EGTA effect on interstitial calcium as GreenT-EC/mCyRFP1 ratio around notochord and dorsal/ventral muscle cells. Regions of interest outlining cell membranes were initially detected automatically using macro-routines and if necessary adjusted manually. **G**) Effects of the CaSR inhibitor Calhex231 applied at 5 µM and 10 µM in E2 embryo medium containing 0.03 mM calcium for 48h. **H**) Effects of Calcitriol (2 µM) treatment for 48 h on interstitial calcium around notochord and muscle. **I**) Effects of large concentration differences of environmental calcium on interstitial calcium in fish larvae. Fish were raised to 2, 4 or 6 dpf (days post fertilization) in E2 embryo medium containing from 0.03 mM to 10 mM calcium and the effects on interstitial calcium were quantified in notochord and muscle tissue. For all experiments each dot represents the mean of n ≥ 6 cells for one fish and each condition included from 3 to 5 fish. Asterisks in figures indicate p values as follows: * p < 0.05, ** p < 0.01, *** p < 0.001.

## Discussion

GreenT-ECs are low affinity calcium sensors with excellent response characteristics. They were tuned to match calcium concentrations found in extracellular body fluids. With the reported properties, they allow detecting relatively small calcium transients in a background of extremely high residual concentrations in free calcium. The sensors were constructed by inserting a small fragment of a calcium binding protein (68 amino acids) into a fluorescent protein. Thus, with appropriate level of engineering, very large fractional fluorescence changes could be obtained without resorting to larger calcium binding proteins, calcium dependent protein interactions or use of circularly permutated fluorescent protein backbones. As common with other fluorescent protein-based single fluorophore sensors, GreenT-ECs, with a pKa of 6.7 (Fig. 2), possess a residual pH-sensitivity in the physiological range. This should be kept in mind and controlled when larger pH changes are expected in a given experimental situation. The indicators were anchored on the surface of expressing cells, resulting in bright fluorescence and high signal-to-noise ratios for imaging. Sparse labeling or cell-type specific labeling also will be possible this way. The alternative would be to have cells secrete GreenT-ECs into the extracellular fluid. However, this might require substantial ubiquitous expression, as sensors might become diluted within body fluids. A secondary application for GreenT-ECs might be its use as a tag to image surface exposure of proteins or secretion, given the bright fluorescence when exposed to high calcium extracellular fluids and dimness when residing in the cytosol (Figure S3).

We validated GreenT-ECs in primary hippocampal neurons. Both in dissociated neurons and in organotypic slices we could show bi-directional fluorescence changes that faithfully reproduced the extracellular decreases or increases in free calcium. We found activity-induced increases in interstitial calcium that appeared driven by cellular extrusion mechanisms. Transient decreases in interstitial calcium due to influx of calcium into the large cytosolic volumes upon neuronal activation were not encountered. GreenT-ECs with their extremely fast off-rate would be very well suited to report such decreases. Possibly the specific distribution of channels, pumps and the dimensions of the intercellular space could contribute to shaping the size and direction of such signals. Dynamics in brain extracellular calcium had been described before^30, 31^. Calcium extrusion from cells was documented using ion selective electrodes in epithelial tissues^32^. Use of membrane-anchored synthetic calcium dyes similarly revealed calcium extrusion in cultured cells *in vitro^33, 34^*.

Transgenic zebrafish lines stably expressing GreenT-EC appear as a promising model of body fluid calcium homeostasis. Using *in vivo* confocal imaging, we could show effects of an inhibitor of the CaSR on uncoupling feed-back loops necessary for maintaining physiological calcium concentrations in body fluids. Similar studies may contribute to elucidate mechanisms and players in the regulation of physiological levels of free calcium in body fluids and their potential compartmentalization across tissues. Dis-regulation of interstitial free calcium is linked to numerous medical disorders in humans^35, 36^. While further investigations into these aspects were outside the scope of this study, these data show the potential of GreenT-ECs for new discoveries in physiology.

## Methods

### Molecular Biology

The plasmid vector pRSET-B (Invitrogen) was used for the library generation, bacterial screenings and protein production. pcDNA 3.1+ (Invitrogen) and pCAGIG (Addgene, USA) were the expression vectors used for the studies in mammalian cells. All constructs were cloned using the homology-based SLiCE methodology^37^. Briefly, fragments were amplified by PCR containing 20 nucleotides overhangs overlapping with the required upstream and downstream sequences. SLiCE extract was prepared as suggested by the authors and 10 µl reactions were used with a vector:insert ratio 1:1. *E. coli* XL1 Blue cells (Invitrogen) were used for all cloning steps and library screenings. Error prone PCR mutagenesis was performed using the GeneMorph II Random mutagenesis kit (Agilent, Germany). Standard PCR reactions for cloning or site-mutagenesis were done using Herculase II Fusion polymerase (Agilent, Germany). All primers were ordered from Eurofins Genomics, Germany.

### Directed evolution

The DNA libraries were incorporated into *E. coli* XL1 Blue cells and transferred onto LB-Agar plates supplemented with ampicillin, obtaining 500-800 colonies per plate. Depending on the size of the library, 20-50 plates were screened in each round. After an overnight incubation at 37 ⁰C, LB-agar plates containing the libraries were further incubated at 4 ⁰C during 24 hours prior imaging to favor protein folding. Using a home-built screening platform^38^, the plates were imaged before and after the addition of 10 ml of screening buffer (30 mM MOPS, 100 mM KCl, 50 mM CaCl_2_, agar 0.5%, pH 7.2) maintained at 42 ⁰C during the screening. Images were acquired and analyzed using a customized python routine^38^. In general, after each round of screening, the selected indicators were expressed and purified for further characterizations. The clones were filtered first according to its response to calcium, then its extinction coefficient, and finally if a suitable candidate was identified, its response was evaluated in mammalian cells. Calibrations and comparisons were made using GCaMP6s as a control indicator. Once an acceptable response to calcium was obtained (> 3000 %), new rounds of screenings were done focusing on improving protein expression, adjusting the kinetic performance and the calcium-binding affinity.

### Protein purification

Proteins with histidine-tags were expressed in *E. coli* BL21 (Invitrogen) overnight at 37 °C in 50 ml auto-inductive LB (LB supplemented with 0.05% D-(+)-glucose (w/v), 0.2% lactose (w/v), 0.6% glycerol (v/v)). Bacteria were harvested by centrifugation (4 °C,10 min, 6000 x g) and re-suspended in 10 ml Resuspension buffer (20 mM Na_2_PO_4_, 300 mM NaCl, 20 mM imidazole) (Sigma Aldrich) supplemented with protease inhibitors (4 μM PMSF, 20 μg/ml Pepstatin A, 4 μg/ml Leupeptin) (Sigma Aldrich), 5 μg/mL DNase and 10 μg/ml RNAse (Sigma Aldrich). Bacteria were first lysed physically through sonication on ice for 7 min 0.8 cycle, 0.8 power output (Bandelin Sonoplus). Insoluble components were pelleted through centrifugation (4 °C, 30 min at 20000 x g). For purification, the supernatant was incubated with 150 μl 6% (v/v) Nickel-IDA agarose bead suspension (Jena Bioscience) for 2 hours at 4 °C under mild agitation. Agarose beads were collected in 1 ml propylene gravity flow columns (Qiagen) and washed with 10 ml Resuspension buffer. The proteins were collected using 700 μl elution buffer (20 mM Na_2_PO_4_, 300 mM NaCl, 300 mM imidazole) (Sigma Aldrich) and dialyzed against MOPS buffer (30 mM MOPS, 100 mM KCl, pH 7.2).

### In vitro spectroscopy

The fluorescence change (ΔF/F_0_) of the indicators was determined in 96-well plates using a fluorescence plate reader (Tecan). The Ca^2+^ free fluorescence (F0) was measured in MOPS buffer supplemented with EGTA 0.4 mM. The fluorescence of the indicator corresponding to the Ca^2+^ bound state was measured in MOPS buffer supplemented with 20 mM Ca^2+^ and 1 mM Mg^2+^.

The molar extinction coefficient (EC) was determined through the absorption of the denatured chromophore at 452 nm (extinction coefficient 44 mM^-1^cm^-1^). Proteins were prepared in MOPS buffer supplemented with 60 mM CaCl_2_ and the absorbance spectrum was acquired before and after the addition of NaOH to a final concentration of 0.025 M. The quantum yield of new variants was determined relative to mNeonGreen by using the slope method. First, the absorption and emission spectra of three serial 1:2 dilutions were acquired in the same cuvette. Then, the integrated emission spectrum was plotted against the maximum absorption and the slope was determined. For mNeonGreen a quantum yield of 0.8 was assumed^19^. The Ca^2+^ affinity of the indicators was determined using MOPS buffer supplemented with 10 mM EGTA, 1 mM Mg^2+^ and increasing concentrations of Ca^2+^ as previously described^39^. Dissociation constant (K_D_) values were determined by plotting the log10 values of the [Ca^2+^] free concentrations in mol/l against the corresponding ΔF/F_0_ values (normalized to ΔF/F_0_ at 39.8 μM Ca^2+^), and fitting a sigmoidal curve to the plot. Prism was used for data analysis.

To determine the pKa of a fluorescent protein (indicating its pH stability), a series of MOPS/MES buffered solutions supplemented with 100 mM Ca^2+^ were prepared, with pH values adjusted in 0.5 pH steps from pH 5.5 to pH 8.5 using NaOH and HCl. In a bottom 96 well plate, triplicates of 200 μl of buffer containing 0.5-1 μM of protein were prepared for each pH value and incubated for 10 minutes at room temperature. Subsequently, all emission spectra were recorded. To determine the pKa value, the relative fluorescence values at the protein’s emission maximum were plotted against the pH values and a sigmoidal fit was applied.

To determine the kinetic rates of the calcium indicators, a Cary Eclipse fluorescence spectrophotometer (Varian) fitted with an RX pneumatic drive unit (Applied Photophysics) was used. For obtaining the macroscopic off-rate constant (Koff), two stock solutions were prepared as follows: a calcium-saturated indicator solution (30 mM MOPS, 50 mM CaCl_2_, 2 mM MgCl2, 100 mM KCl, ∼ 0.2–1 μM indicator, pH 7.2) and a BAPTA solution (30 mM MOPS, 100 mM KCl, 100 mM BAPTA, pH 7.2). The stopped-flow experiment was carried out at room temperature (∼23 °C) and the two solutions were mixed with an injection pressure of 3.5 bar. Excitation was set to 480 nm and emission was detected at 520 nm. The acquisition time was set to 12.5 ms, duration to >10 s and mixing volume to 400 μl with a mixing dead time of the instrument of 8 ms. The decay time (τ, s) was determined by fitting with a double-exponential curve to the fluorescence response using Prism. Macroscopic on-rate kinetics (Kobs) were obtained by mixing the calcium-free buffer containing the protein (30 mM MOPS, 100 mM KCl, 1 mM MgCl_2_, ∼ 0.2–1 μM indicator, pH 7.2) and solutions containing increasing concentrations of CaCl_2_ (30 mM MOPS, 100 mM KCl, 0.1-100 mM CaCl_2_, 1 mM MgCl_2_, ∼ 0.2–1 μM indicator, pH 7.2). Concentrations of free calcium were calculated using WEBMAXC STANDARD.

### Crystallization and Structure Determination

GreenT-EC crystals were formed using the sitting drop method. The precipitant solution was 0.2 M ammonium sulfate with 30% w/v PEG 400. Droplet conditions were: 0.4 µL total volume, 200 nL protein + 200 nL precipitant. The crystals were cryoprotected with addition of 30% of ethylene glycol and flash frozen in liquid nitrogen. X-ray data sets were recorded on the 10SA (PX II) beamline at the Paul Scherrer Institute (Villigen, Switzerland) at wavelength of 1.0 Å using a Dectris Eiger 16M detector with the crystals maintained at 100K by a cryocooler. Diffraction data were integrated using XDS (BUILT=20220220) and scaled and merged using AIMLESS (0.7.7); data collection statistics are summarized in Supplementary Table 2. Initially the GreenT-EC data set was automatically processed at the beamline to 1.3 Å resolution and a structure solution was automatically obtained by molecular replacement using pdb 5MWC as template. The map was of sufficient quality to enable 90 % of the residues expected for GreenT-EC to be automatically fitted using Phenix autobuild. The model was finalised by manual rebuilding in COOT2 (0.9.6) and refined using in Phenix refine (1.19.2).

### Cell lines and Tissue Culture

HeLa and HEK 293T (Invitrogen) cells were grown in high glucose Dulbecco’s Modified Eagle Medium with high glucose, pyruvate (Gibco) supplemented with 10% fetal bovine serum (FBS), 29.2 mg/ml of L-glutamine, 10,000 units of penicillin and 10,000 µg streptomycin at 37°C with 5% CO_2_.

### Evaluation of indicator surface display

Different combinations of signal peptides and membrane domains were evaluated in HeLa cells. The cells were seeded in 35 mm glass-bottom dishes (Mat-Tek) pre-coated with Poly-L-lysine (Sigma) and transfected with the different constructs one day before imaging, according to the manufacturer instructions (Lipofectamine 3000). Hoechst staining was performed immediately before imaging using a final concentration of 2 µg/mL for 20 minutes. The imaging buffer used for these experiments was Hank Balanced Salt Solution (HBSS) supplemented with 1 mM MgCl_2_ and 3 mM CaCl_2_. Images were acquired in a Leica Stellaris5 microscope with a 60x oil-immersed objective. Different export signal peptides and transmembrane domain were evaluated: IβCE (from Integrin-β of C. elegans)^40, 41^, Igk (from human CH29 light chain)^40, 42^, Nrxn1 (mouse Neurexin I first 63 aa)^40, 42^, PPA (N-terminal 24 aa of mouse preproacrosin signal peptide)^24^, PDGFRB (Homo sapiens platelet derived growth factor receptor beta)^43^, GPI (mouse Thy-1 glycosylphosphatidylinositol) anchoring domain^24^, NXN (rat neurexin-1β) and NLG (rat neuroligin-1)^40, 42^.

### Evaluation of reference proteins using Igk-PDGFR surface localized GreenT-EC

HeLa cells were seeded in 35 mm glass-bottom dishes (Mat-Tek) and transfected with the different constructs according to the manufacturer instructions (Lipofectamine 3000). The imaging buffer used for these experiments was Hank Balanced Salt Solution (HBSS) supplemented with 1 mM MgCl_2_ and 3 mM CaCl_2_. Images were acquired in a Nikon Spinning Disk confocal microscope with a 20x oil-immersed objective. In each case, the reference proteins mCarmine^38^, mScarlet^44^, mCyRFP1^23^, mCerulean3^45^ and mTurquoise2^46^ were inserted in the C-terminal extreme of the PDGFR domain, separated by a short linker.

### Affinity titrations on the surface of HEK 293T cells

HEK293T cells were seeded on day one into 35 mm glass-bottom dishes (Matek) pre-coated with Poly-L-lysine (Sigma). On day three, the cells were transfected using 1 µg of the plasmids coding for the different variants according to the manufacturer instructions (Lipofectamine 3000). For each variant, three dishes were prepared. On day four, cells were washed with buffer MOPS 30 mM KCl 100 mM, EGTA 0.4 mM, pH 7.2 during 2 min. Then 3 ml of buffer MOPS 30 mM KCl 100 mM, pH 7.2 was added and cells were imaged in a Leica SP8 confocal microscope. For the titration, 80-100 µl of CaCl_2_ stock solutions were added drop-wise to the dishes during time-lapsed imaging. Normally, images were acquired every 1 min. All experiments were done in a SP8 confocal microscope (Leica) using a 63x water-immersed objective.

### Image processing and ratioing

The general processing procedure for mammalian cell titrations and zebrafish experiments consisted in: i) Noise reduction (typically using a median or Gaussian filter of 2 pixels), ii) Thresholding saturated pixels and noise/cytoplasm signals. During this process, the original image was divided by the Threshold mask to assign N/A values to the pixels that were thresholded out. This avoids problems arising from averaging cero-value pixels during the ROI analysis. iii) Finally, the processed GreenT-EC and mCyRFP1 channels were used to obtain a ratiometric image GreenT-EC/mCyRFP1. This method allowed us to monitor the ratio signals in the membrane with minor interference from background or low intensity cytosolic signals. ImageJ macro routines were generated to perform the image processing, ROI detection/drawing, measurement of individual channels and ratio, and exporting of the results. The data analysis was done in GraphPad. Images were displayed with white background in cases where high contrast was required to clearly visualize the different structures.^47^

### Primary neuronal cell culture

All experiments were performed in accordance with the European directive on the protection of animals used for scientific purposes (2010/63/EU). Banker culture of hippocampal neurons were prepared from 18 day pregnant embryonic Sprague-Dawley rats as described previously ^48^. Briefly, hippocampi were dissected in HBSS containing Penicillin-Streptomycin (PS) and HEPES and dissociated with Trypsin-EDTA/PS/HEPES. Neurons were plated in minimum essential medium supplemented with 10% horse serum on coverslips coated with 1 mg/ml poly-L-lysine in 60 mm petri dishes at a density of 250000 cells per dish. Following neuronal attachment, the coverslips were flipped onto 60-mm dishes containing a glial cell layer in Neurobasal medium supplemented with L-glutamine (GIBCO, #25030-024) and NeuroCult SM1 Neuronal Supplement (StemCell Technologies, #05711). Cells were maintained at 37°C with 5% CO_2_ for 13-15 days. Neurons were transfected at DIV 7-9 using the calcium-phosphate co-precipitation method. Per dish, precipitates containing 12 µg plasmidic DNA of GreenT-EC-GRAPHIC were prepared using the following solutions: TE (1mM Tris–HCl pH 7.3, 250 mM EDTA pH 8), CaCl_2_ (2.5 mM CaCl_2_ in 10 mM HEPES, pH 7.2), 2 x HEPES-buffered saline (HEBS; 11 mM D-Glucose, 42 mM HEPES, 10 mM KCl, 270 mM NaCl and 1.4 mM Na_2_HPO_4_.2H_2_O, pH 7.2). Coverslips containing neurons were moved to 12 well multi-well plates containing 250 ml/well of conditioned culture medium. The 50 µl precipitate solution was added to each well, in the presence of 2 mM kynurenic acid (Sigma-Aldrich #K3375) and incubated for 1h-1h30 at 37°C. Then, cells were washed with non-supplemented Neurobasal medium containing 2 mM kynurenic acid for 20 min and moved back to their original culture dish. Cells were imaged at DIV 13-15.

### Organotypic hippocampal slice culture

Mice were housed under a 12 h light/12 h dark cycle at 20-22 °C with ad libitum access to food and water in the animal facility of the Interdisciplinary Institute for Neuroscience (University of Bordeaux/CNRS), and monitored daily by trained staff. All animals used were free of any disease or infection at the time of experiments. Pregnant females and females with litters were kept in cages with one male. We did not distinguish between males and females among the perinatal pups used for organotypic cultures, as potential anatomical and/or physiological differences between the two sexes were considered irrelevant in the context of this study. C57Bl/6J wild-type mice were used for all brain slice experiments in this study. Brain organotypic cultures were prepared according to the method described previously^49^. Briefly, hippocampal slices were obtained from postnatal P5-7 old C57BI/6J mouse pups. The animals were quickly decapitated and the brains placed on cold sterile dissection medium (all in mM; 0.5 CaCl_2_, 2.5 KCl, 2 MgCl_2_, 0.66 KH_2_PO_4_, 0.85 Na_2_HPO_4_-12H_2_O, 0.28 MgSO_4_-7H_2_O, 50 NaCl, 2.7 NaHCO_3_, 25 glucose, 175 sucrose, 2 HEPES; all from Sigma). The hippocampi were dissected and sliced on a McIlwain tissue chopper to generate coronal brain slices of 350 mm thickness. After 20 minutes of incubation at 4°C, the slices were transferred on sterilized hydrophilic polytetrafluoroethylene (PTFE) membrane (FHLC04700; Merck Millipore) pieces, which were placed on top of cell culture inserts (Millipore, 0.4 mm). The inserts were held in a 6-well plate filled with medium (50% Basal medium eagle (BME), 25% Hank’s Balanced Salt Solution (HBSS, pH 7.2), 25% Horse Serum, 11.2 mmol/L glucose and 20 mM glutamine; all from GIBCO) and cultured up to 14 days at 35°C/5% CO_2_. Culture medium was replaced every two days.

### AAV production and injections

In order to introduce the GreenT-EC plasmid to neurons in organotypic brain slices, the GreenT-EC-GRAPHIC coding sequence was first inserted into a specific viral plasmid (Addgene #50477) with a neuron-specific promotor CaMKII. After testing the plasmid for its correct size and integrity via restriction enzyme digestion, the construct was sent to in-house virus production facility, which provided us with a construct of GreenT-EC-GRAPHIC integrated into AAV9 particles. The viral particles were injected into the hippocampal slices via microinjections using a glass pipette connected to Picospritzer (Parker Hannifin). Briefly, the virus was injected via a pipette positioned into the CA1 area of the slice by brief pressure pulses (30 ms; 15 psi). The virus was injected at least 7 days prior to the experiments.

### STED and confocal microscopy of hippocampal neurons

We used a custom-built 3D-STED/confocal setup^50^ constructed around an inverted microscope body (DMI 6000 CS, Leica Microsystems) which was equipped with a TIRF oil objective (x100, 1.47 NA, HXC APO, Leica Microsystems) and a heating box (Cube and Box, Life Imaging Services) to maintain a stable temperature of 32°C. A pulsed-laser (PDL 800-D, PicoQuant) was used to deliver excitation pulses at 488 nm and a de-excitation laser (Onefive Katana 06 HP, NKT Photonics) operating at 594 nm was used to generate the STED light pulses. The STED beam was profiled to a donut shape using a spatial light modulator (Easy3D Module, Abberior Instruments). Image acquisition was controlled by the Imspector software (Abberior Instruments). The spatial resolution for the excitation beam was 175 nm (x-y) and 450 nm (z) and for the STED beam 60 nm (x-y) and 160 nm (z).

### Image acquisition of hippocampal neurons

For imaging, either slices or dissociated neurons were transferred on their glass coverslip to an imaging chamber and immersed in an imaging medium (in case of slices: artificial cerebrospinal fluid, ACSF; consisted of (in mM) 119 NaCl, 2.5 KCl, 1.3 MgSO_4_, 1 NaH_2_PO_4_ x 2H2O, 1.5 CaCl_2_* x 2H_2_O, 20 D-Glucose x H_2_O and 10 HEPES (all from Sigma Aldrich); 300 mOsm; pH 7.4; in case of dissociated neurons: Tyrode solution; consisted of (in mM) 10 D-Glucose x H_2_O, 100 NaCl, 5 KCl, 2 MgCl_2_ x 6H_2_O, 25 HEPES, 2 CaCl_2_* x 2H_2_O). Confocal time-lapse images of dissociated neurons had a field of view: 50 µm x 50 µm. STED single-plane images of organotypic brain slices were either 100 µm x 100 µm or 50 µm x 50 µm with a pixel size of 48,6 nm, 0.3 ms dwell time. The excitation power was 0.5 µW (measured before the objective) and the STED power was 30 mW. *The concentration of calcium in the solutions varied depending on the experiment.

### 2-photon microscopy

We used a commercial two-photon microscope (Prairie Technologies) and 40X water immersion objective (NA 1.0; Plan-Apochromat, Zeiss). For GreenT-EC excitation, a two-photon laser (TI:sapphire, Mai Tai, Spectra Physics) was tuned to 900 nm; with laser power ranging from 10-25 mW in the focal plane. The fluorescence signal was collected in a non-descanned manner by PMT detectors. The imaging parameters were adjusted using the commercial software provided by Prairie. Image acquisition Individual slices were transferred to a submerged recording chamber and continuously bathed with HEPES-based ACSF solution (in mM): 119 NaCl, 2.5 KCl, 1.3 MgSO_4_, 1 NaH_2_PO_4_ x 2H_2_O, 20 D-Glucose x H_2_O, 10 HEPES and varying CaCl_2_ concentrations: 0, 1.5, 2.5, 8, 15 (all from Sigma Aldrich); 300 mOsm; pH 7.4. For most of the experiments, we acquired time-lapse images of single planes (126 µm x 126 µm) every 1 s for a total number of 40 repetitions. If the imaging parameters differed, it is mentioned in the figure legend related to a specific part of the figure.

### Calcium puffing experiments

In order to locally apply calcium solutions, we inserted a glass micropipette into the slice expressing GreenT-EC sensor. By injecting brief high-pressure pulses (10-100 ms, 15 psi) via a Picospritzer (Parker Hannifin), calcium solution was locally delivered into the desired imaging region. To avoid pressure-induced z-drift the injection parameters, pulse durations, pressure levels, were optimized.

### Electrophysiology

Schaffer collateral fibers in hippocampal slices were electrically stimulated and evoked field excitatory postsynaptic potentials (fEPSP) were recorded in the Stratum radiatum of hippocampal CA1. Two glass micro-electrodes (tip resistance 5-6 MΩ) for stimulation and recording were filled with aCSF and carefully positioned in the slice and placed at depths where imaging was performed. The current pulses (5-15 pulses, 0.2-0.3 ms in duration) were delivered via the stimulating electrode from a stimulus isolator (AMPI; Science Products). The stimulus strength varied between 10-40 µA. The field potentials were recorded using a patch clamp amplifier (Multiclamp 700B; Molecular Devices).

To block calcium extrusion from cells, we applied two calcium-pump blockers: 5 µM sodium-orthovanadate (Sigma Aldrich) and 50 µM benzamil hydrochloride hydrate (Sigma Aldrich) dissolved in HEPES-based aCSF.

### Data analysis and statistics of brain slices

All images were processed and analyzed using the ImageJ software. Statistical tests were performed using Graphpad Prism software. Normally distributed data are presented as mean with standard deviation, while non-normal data are presented as median with interquartile range. The size and type of individual samples, n, for given experiments is indicated and specified in the results section and in figure legends. Asterisks in figures indicate p values as follows: * p < 0.05, ** p < 0.01, *** p < 0.001.

Normal distribution was tested Shapiro-Wilk test. One-way ANOVA followed by a Dunnett’s test was used to evaluate differences among control and treated groups.

### Zebrafish strains and husbandry

Transgenic zebrafish derived from the outbred AB strain were used in all experiments. Zebrafish were raised under standard conditions at 28 °C. Animals were chosen at random for all experiments. Zebrafish husbandry and experiments with all transgenic lines were performed under standard conditions as per the Federation of European Laboratory Animal Science Associations (FELASA) guidelines^51^, and in accordance with institutional (Université Libre de Bruxelles (ULB)) and national ethical and animal welfare guidelines and regulation, which were approved by the ethical committee for animal welfare (CEBEA) from the Université Libre de Bruxelles (protocols 578N-579N).To generate the actb2:GreenT-EC construct, the vector containing 9.8 kb of zebrafish b-actin2 (*actb2*) promoter^25^ was digested by SpeI/NotI and GreenT-EC cloned using the previously mentioned SLiCE methodology^37^. Transgenics were generated using the I-SceI meganuclease system. Two founders were isolated and screened for the strength of green and red fluorescence in the body. The line with brighter expression was used to perform all experiments, and was designated *Tg(actb2:GreenT-EC)^ulb18^*.

For experiments, transgenic zebrafish larvae, *Tg(actb2:GreenT-EC),* were obtained from outcross of the transgenic adult animal to AB wild-type animals. Larvae between 2 and 6 dpf (days post-fertilization) were used in all experiments. Zebrafish have indeterminate growth with the rate varying with parameters such as fish density, so larvae were kept at a density of 1 larvae per 1ml of E2 medium (7.5 mM NaCl, 0.25 mM KCl, 0.5mM MgSO_4_, 75 mM KH_2_PO_4_, 25 mM Na2HPO_4_, 0.35 mM NaHCO_3_) supplemented with either 0.3, 0.03, 2 or 10 mM CaCl_2_, 0.5 mg/L methylene blue, pH 7.4 since birth unless specified otherwise. Anaesthesia was administered in E2 medium using 0.02% pH 7.0 tricaine methanesulfonate (MS-222; E10521; Sigma-Aldrich, Darmstadt, Germany).

### Zebrafish confocal image acquisitio**n**

Animals were anesthetized in 0.02 % tricaine methanesulfonate (MS-222; E10521; Sigma-Aldrich, Darmstadt, Germany) and mounted in 1% Low-Melt Agarose (50080; Lonza, Basel, Switzerland) and imaged on a glass-bottomed FluoroDish (FD3510-100; World Precision Instruments (WPI), Sarasota, Florida) using a LSM 780 confocal microscope (Zeiss). Finfold and muscle were imaged using a 40x/1.1N.A. water correction lens. Imaging frame was set at 1024 x 1024, and the distance between confocal planes was set up at 3 µm for Z-stack cover, on average, a thickness of 60 µm. Samples were excited with 488 nm laser and fluorescence was collected in the two channels simultaneously using detector width of 493-530 nm for GreenT-EC and 550-740 nm for mCyRFP1.

### Zebrafish pharmacological treatments

All compounds for treating embryos were dissolved in DMSO according to manufacturer, diluted in E2 embryo medium (0.3 mM or 0.03 mM Ca^2+^), changed daily and embryos treated by immersion. The compounds, and concentrations used, with catalogue numbers were: EGTA (Roth), 0.3 mM for 10 min; Calhex 231 hydrochloride (SML0668-Sigma), 5 or 10 µM for 48h (from 2 to 4 dpf); Calcitriol (C3078-TCI) 2.5 µM for 24h (3 to 4dpf).

### Zebra fish data analysis and statistics

All images were ratiometrically processed as described and analyzed using the ImageJ software with a custom-made macro (available upon request). For each fish, n ≥ 6 cells/region of interested were analyzed for the different tissues (Notochord, muscle or epithelium). The mean ratio GreenT-EC/mCyRFP1 was then plotted for each animal. All experiments included between 3 to 5 animals per group. Statistical analyses were performed using Graphpad Prism software. Data was tested for normality using the Saphiro-Wilk test. Normally distributed data is presented as mean with standard deviation and was analyzed using 2-way ANOVA followed by a Tukey’s or Šídák’s multiple comparisons test to evaluate differenced among control and treated groups in different tissues/regions of interest. The size, n, for given experiments are indicated in figure legends. Asterisks in figures indicate p values as follows: * p < 0.05, ** p < 0.01, *** p < 0.001.

### Availability of Material

Mammalian and bacterial expression vectors coding for GreenT-ECs will be made available from Addgene. The crystal structure of variant GreenT NRS F241Y will be deposited at https://www.rcsb.org/

## Acknowledgements

We thank the Protein Core Facility of the Max-Planck-Institute for Biochemistry for their assistance. This work was supported by the Max-Planck-Society, by MISU funding from the FNRS (34772792, 40005588) and from Fondation Jaumotte-Demoulin to S.P.S. and I.G.

## Author contributions

O.G. led the study. AVG, VN, SS and OG conceived experiments. AVG conducted protein engineering, directed evolution, in vitro spectroscopy, titrations, microscopy of cell lines, image processing. AF conducted protein evolution screenings and developed python scripts for data acquisition and analysis. IGS generated transgenic zebrafish and performed in vivo confocal experiments on fish, AI performed experiments in rodent hippocampus, JA participated in neuronal data collection, AI and VN analyzed experiments in rodent hippocampus, JB crystallized GreenT-EC NRS F241Y, AVG and OG wrote the manuscript, all authors contributed to discussing and revising the manuscript.

## Supplementary Figure legends

**Figure S1:**
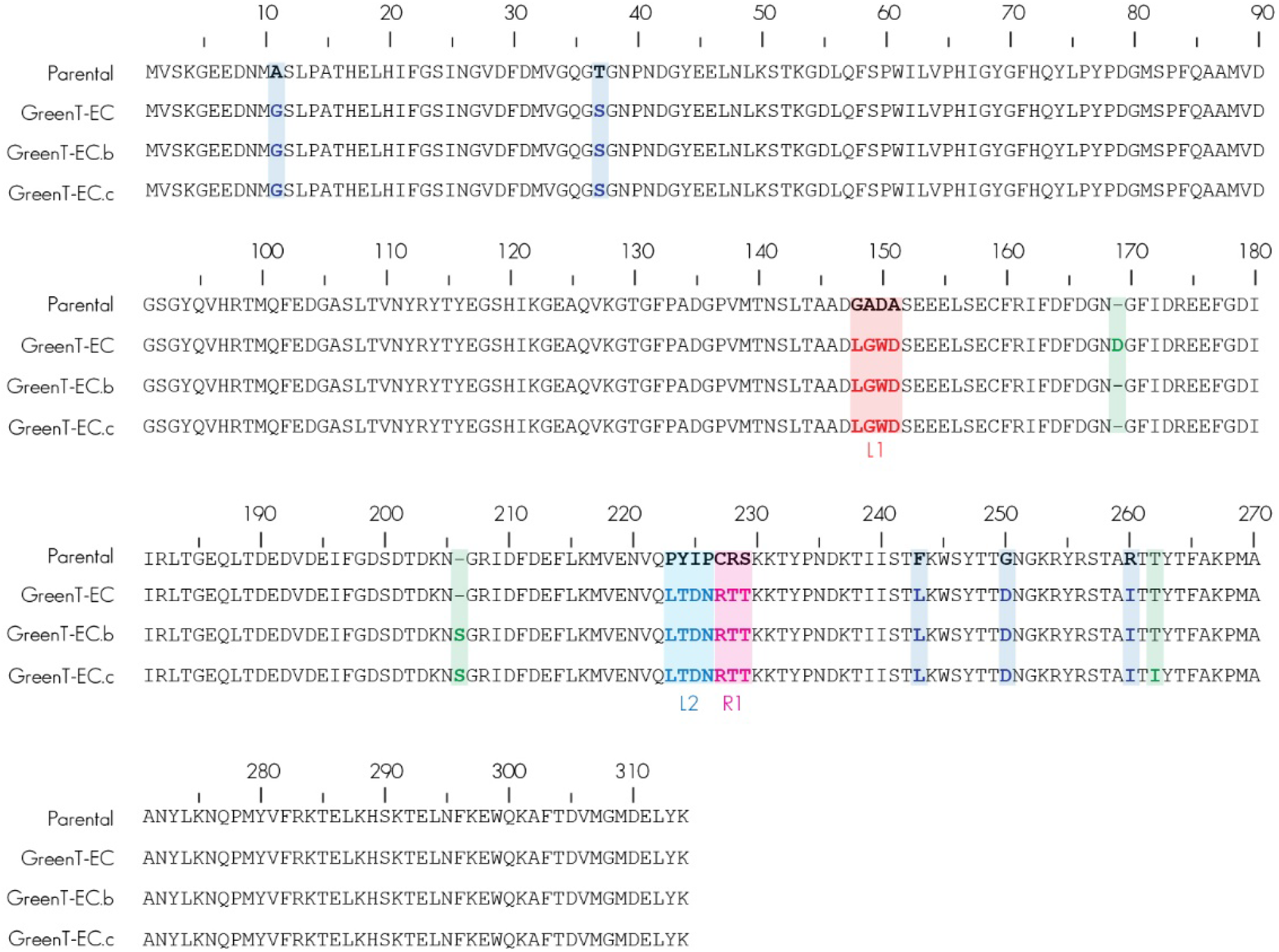
Amino acid sequences of GreenT-ECs and the parental protein. The amino acid sequence of the parental protein that served as starting point for directed evolution of GreenT-ECs is shown in the top row. While GreenT-EC contains an insertion of an aspartic acid after residue 169 in the in the first calcium binding EF-hand motif (colored green), GreenT-EC.b and GreenT-EC.c contain insertions of a serine (colored green) after position 205 within the second calcium binding EF-hand motif. GreenT-ECc contains an additional amono acid exchange (T260I, green). Engineered residues of linker 1 (L1) between mNeonGreen and the minimal TnC domain are in red. Residues marked in blue represent the engineered linker 2 (L2) between mNeonGreen and the minimal calcium binding domain. Residues highlighted in magenta adjacent to the TnC domain correspond to region R1. Amino acid exchanges in mNeonGreen that enhanced response, folding and expression of GreenT-ECs are colored dark blue. Numbering is based on the parental construct. The insertion of D169 or S205 will shift it accordingly.

**Figure S2:**
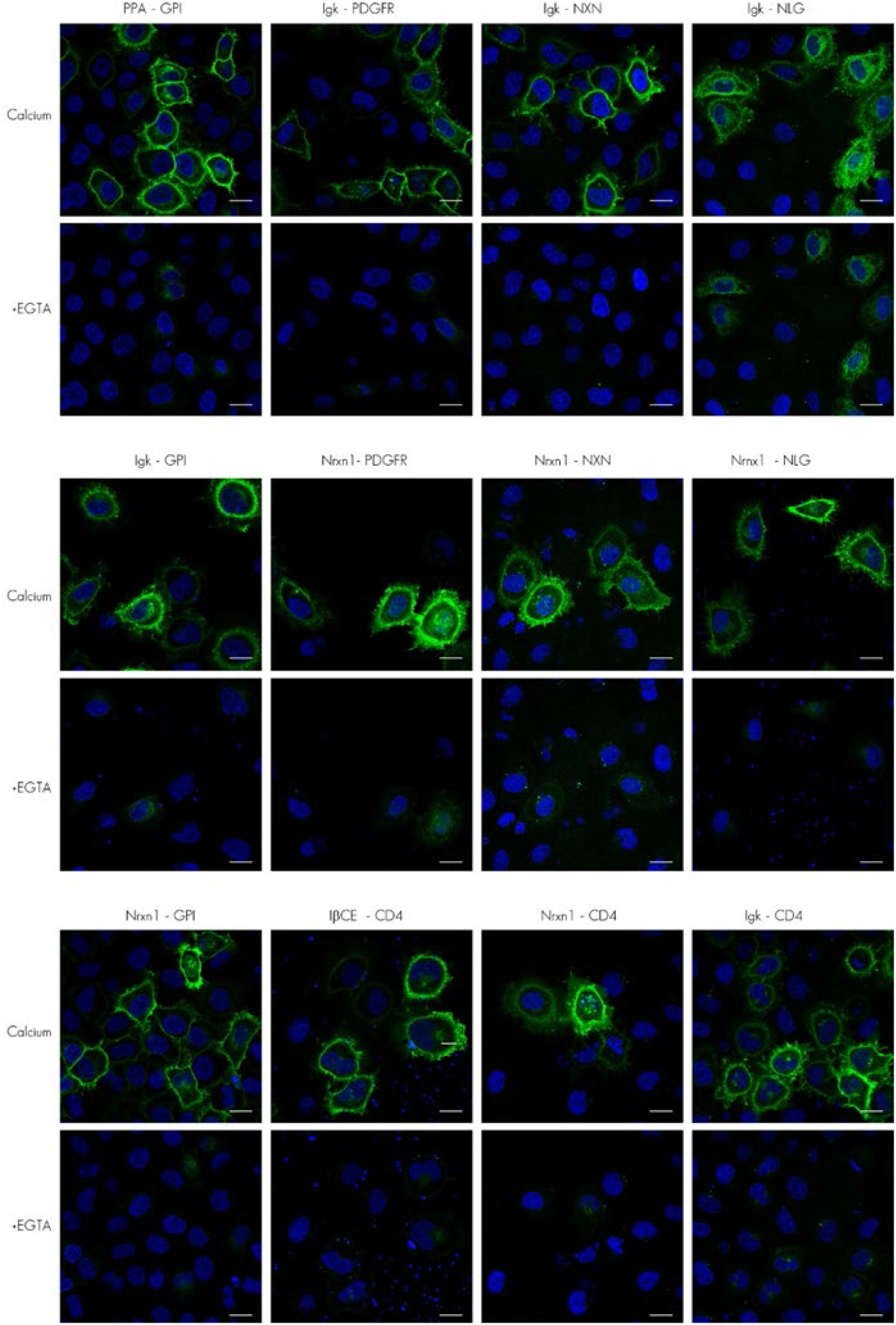
Surface delivery of GreenT-ECs. Exemplary images show expression of constructs in which different surface targeting sequences were fused to GreenT-EC. Cells are shown with either 3 mM extracellular calcium concentration in HBSS buffer or at zero extracellular calcium using 3 mM EGTA. GreenT-EC fluorescence (green) and Hoechst 33342 nuclear counterstain (blue) are displayed. Scale bar, 20 µm. CD4: transmembrane domain of the T-cell surface glycoprotein Cluster of Differentiation 4. PPA: N-terminal 24 amino acids of mouse preproacrosin signal peptide. PDGFR: Homo sapiens platelet derived growth factor receptor beta; NXN: rat neurexin-1β; NLG: rat neuroligin-1. GPI: mouse Thy-1 glycosylphosphatidylinositol anchoring domain; Igk: N-terminal 21 aa from human CH29 light chain; Nrxn1: N-terminal 63 aa of mouse Neurexin I; IßCE: N-terminal 29 amino acids from Integrin-ß of C. elegans.

**Figure S3:**
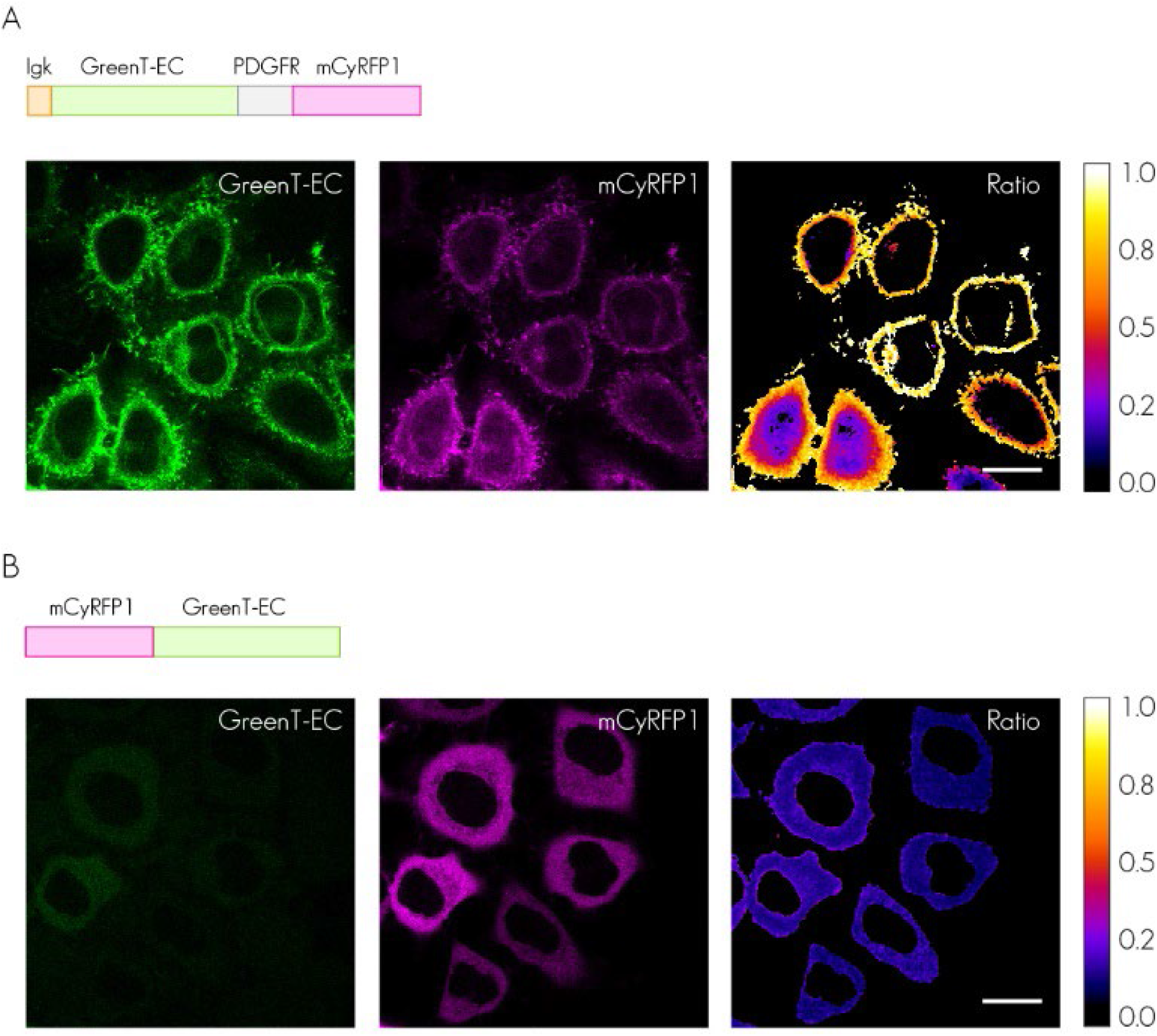
Cytosol vs Surface Localization of GreenT-EC in HEK293 cells. **A)** GreenT-EC fluoresces brightly when cell surface localized and exposed to high calcium extracellular buffer. Green channel: GreenT-EC fluorescence. Red channel: mCyRFP1 reference protein. Ratio: GreenT-EC/mCyRFP1. **B**) GreenT-EC is essentially non-fluorescent when localized within the cytosol. Scale bar, 50 µm.

**Figure S4:**
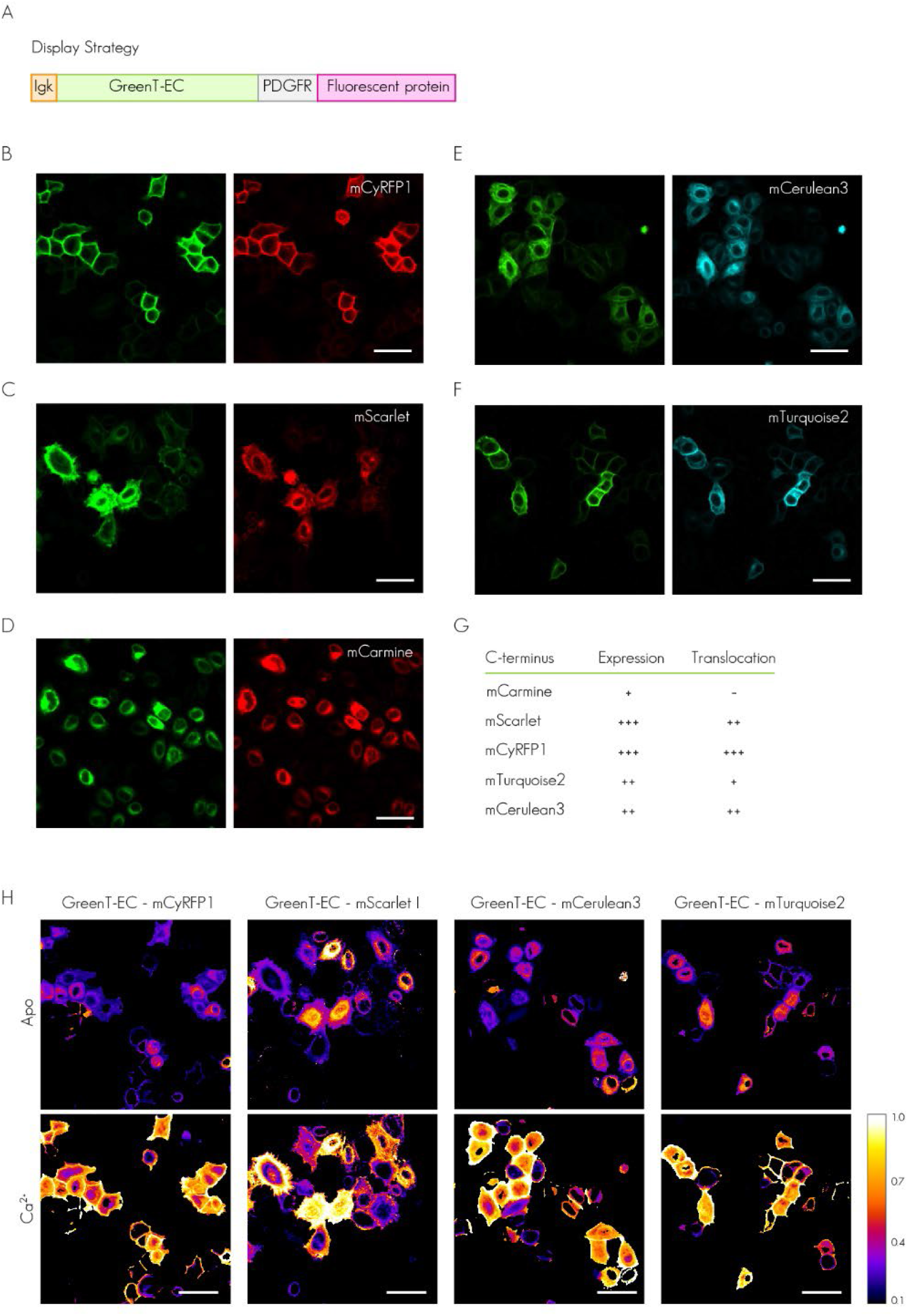
Fusions of reference proteins to the cytosolic tail of GreenT-EC. **A**) Schematic representation of the surface-targeted GreenT-EC construct used for evaluation. It consists of the export signal peptide (Igk), the sensor (GreenT-EC), the transmembrane domain (PDGFR) and the cytosolic fusion of the reference proteins. The use of mCyRFP1 **(B)**, mScarlet-I **(C)**, mCarmine **(D)**, mCerulean3 **(E)** and mTurquoise2 **(F)** was evaluated in HeLa cells in the presence of 3 mM Ca^2+^ and its performance was qualitatively rated in terms of expression and membrane/cytosol contrast (translocation). While mScarlet, mCyRFP1, mCerulean3 and mTuquoise2 showed a clear contrast membrane/cytosol, mCarmine was only visible in the cytosol indicating that is accumulated inside cells during protein translocation. **G)** The information was qualitatively summarized in the table. **H)** Ratiometric images GreenT-EC/Reference protein were calculated in the presence of Ca^2+^ 3 mM before and after the addition of EGTA 3 mM. Scale bar, 50 µm.

**Figure S5:**
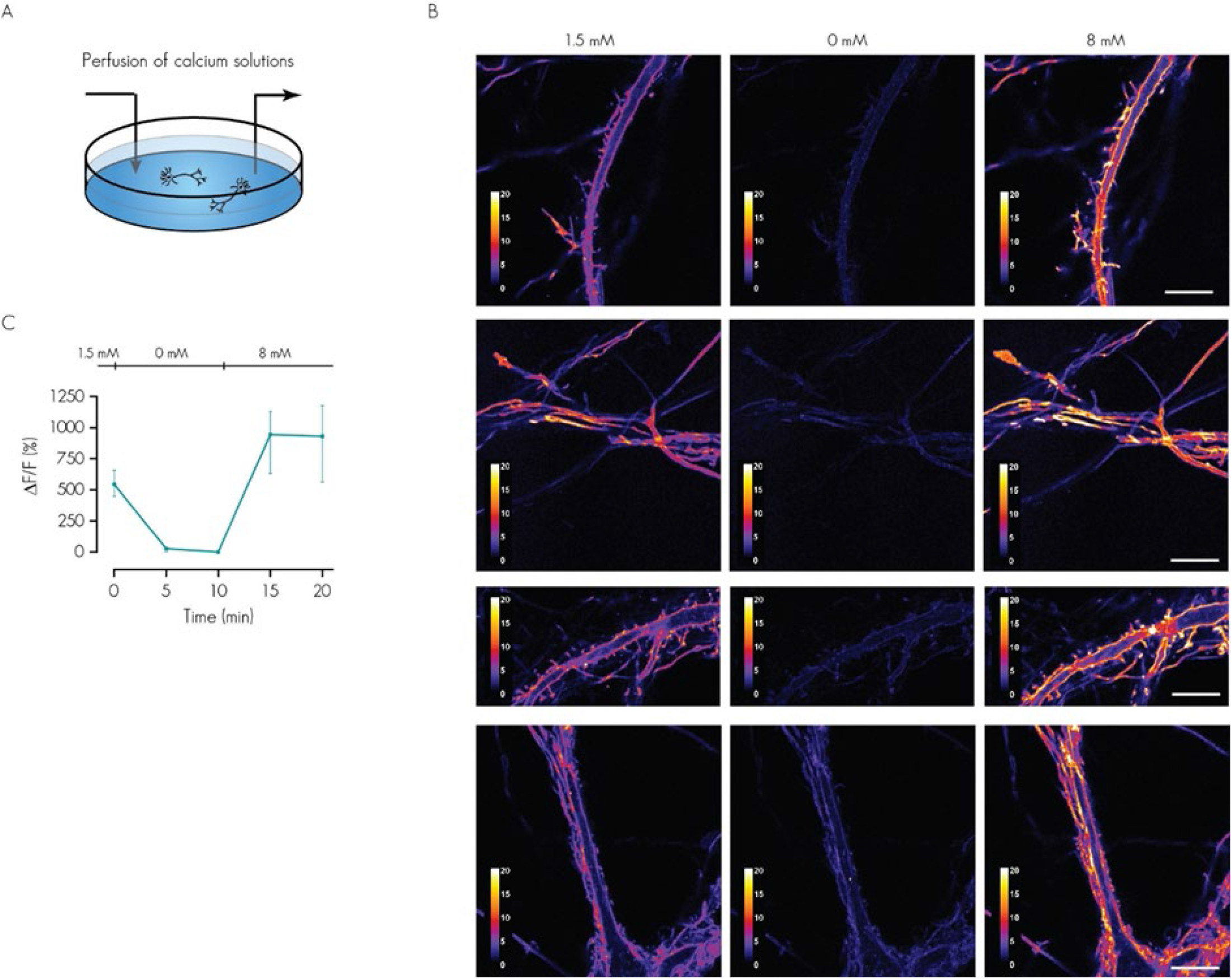
GreenT-EC fluorescence intensity changes in primary hippocampal neurons in response to different extracellular calcium concentrations. **A)** Schematic representation of perfusion experiments. **B)** Confocal time-lapse imaging. Images used to quantify fluorescence changes are displayed using a pseudo-color LUT to facilitate visualization of the extremely dim 0 mM calcium condition. Scale bar, 10 µm. Quantification of fluorescence changes compared to 0 mM point respectively during the perfusion experiments. Error bars are standard deviations and correspond to three independent neuronal culture experiments.

---

**Table S1.** Crystal structure: Data collection and refinement statistics.

|  | **GreenT-EC\*** |
| --- | --- |
| **Wavelength** | 1.0 |
| **Resolution range** | 47.85 - 1.279 (1.325 - 1.279) |
| **Space group** | P 21 21 21 |
| **Unit cell** | 54.332 62.854 95.709 90 90 90 |
| **Total reflections** | 1047921 (90739) |
| **Unique reflections** | 85238 (8382) |
| **Multiplicity** | 12.3 (10.8) |
| **Completeness (%)** | 99.97 (99.73) |
| **Mean I/sigma(I)** | 15.60 (0.94) |
| **Wilson B-factor** | 15.85 |
| **R-merge** | 0.08807 (2.069) |
| **R-meas** | 0.09188 (2.171) |
| **R-pim** | 0.02593 (0.6485) |
| **CC1/2** | 0.999 (0.389) |
| **CC\*** | 1 (0.748) |
| **Reflections used in refinement** | 85233 (8382) |
| **Reflections used for R-free** | 4303 (405) |
| **R-work** | 0.1832 (0.3259) |
| **R-free** | 0.2005 (0.3231) |
| **CC(work)** | 0.962 (0.671) |
| **CC(free)** | 0.967 (0.672) |
| **Number of non-hydrogen atoms** | 2624 |
| **macromolecules** | 2249 |
| **ligands** | 44 |
| **solvent** | 343 |
| **Protein residues** | 282 |
| **RMS(bonds)** | 0.010 |
| **RMS(angles)** | 1.11 |
| **Ramachandran favored (%)** | 99.64 |
| **Ramachandran allowed (%)** | 0.36 |
| **Ramachandran outliers (%)** | 0.00 |
| **Rotamer outliers (%)** | 0.41 |
| **Clashscore** | 7.05 |
| **Average B-factor** | 21.46 |
| **macromolecules** | 20.10 |
| **ligands** | 23.85 |
| **solvent** | 30.16 |

Statistics for the highest-resolution shell are shown in parentheses.

